# Spx inhibits expression of the SwrA•DegU master flagellar activator in *Bacillus subtilis*

**DOI:** 10.1101/2025.09.12.675978

**Authors:** Abigail E. Jackson, Stephen G. Olney, Daniel B. Kearns

## Abstract

Flagella are trans-envelope nanomachines expressed from genes organized in a complex regulatory hierarchy governed at the highest level by transcription factors called master activators. The master activator of flagellar biosynthesis in Bacillus subtilis is a hybrid of SwrA•DegU that is required to increase flagellar density to swarm over solid surfaces. Here we show that the ClpX unfoldase subunit of the ClpP protease is required for swarming motility, and that cells mutated for ClpX fail to swarm due to restricted levels of both SwrA and DegU. Suppressor mutations were found that increased expression of the *fla/che* operon under SwrA•DegU control, and mutation of the LonA protease elevates the levels of SwrA protein while mutation of the global transcriptional regulator Spx increases transcription of both the *degU* and *swrA* genes. We conclude that ClpX promotes swarming motility via degradation of Spx, which represses motility gene transcription including the P*_fla/che_*, P*_degU_*, and P*_swrA_* promoters each activated by DegU. The ClpX-dependent regulatory proteolysis of Spx is relieved upon stress conditions, and we infer that Spx may dampen DegU-mediated positive feedback to limit cell envelope stress caused by excessive flagellar biosynthesis.

## INTRODUCTION

Ancestral strains of *Bacillus subtilis* exhibit two forms of flagellar-mediated motility, called swimming and swarming (Mukherjee 2014). Swimming motility occurs in a liquid environment and involves the movement of individuals in three dimensions, while swarming motility occurs on a solid surface and cells move as groups in two dimensions (Kearns 2010). While both swimming and swarming require the same flagellar structural genes, swarming has additional genetic requirements. For example, swarming requires the production of surfactin, a secreted surfactant that reduces surface tension and creates a thin layer of water within which to swarm (Kearns 2003; Julkowska 2005). Swarming cells are also hyperflagellated compared to swimming cells and require enhanced activity of the SwrA•DegU hybrid master regulator of flagellar biosynthesis to increase flagellar gene expression (**Fig 1**) (Kearns 2005; Calvio 2005; Mukherjee 2015; Mishra 2004). Commonly-used domesticated laboratory strains lack swarming motility due to mutations in *sfp* and *swrA* that abolish surfactin production and hyperflagellation, respectively (Nakano 1992, Kearns 2003, Kearns 2004, Julkowska 2005; Calvio 2005). To better understand swarming motility, we identify and characterize other genes that alter swarming behavior in the ancestral strain of *B. subtilis*.

**Figure 1:**
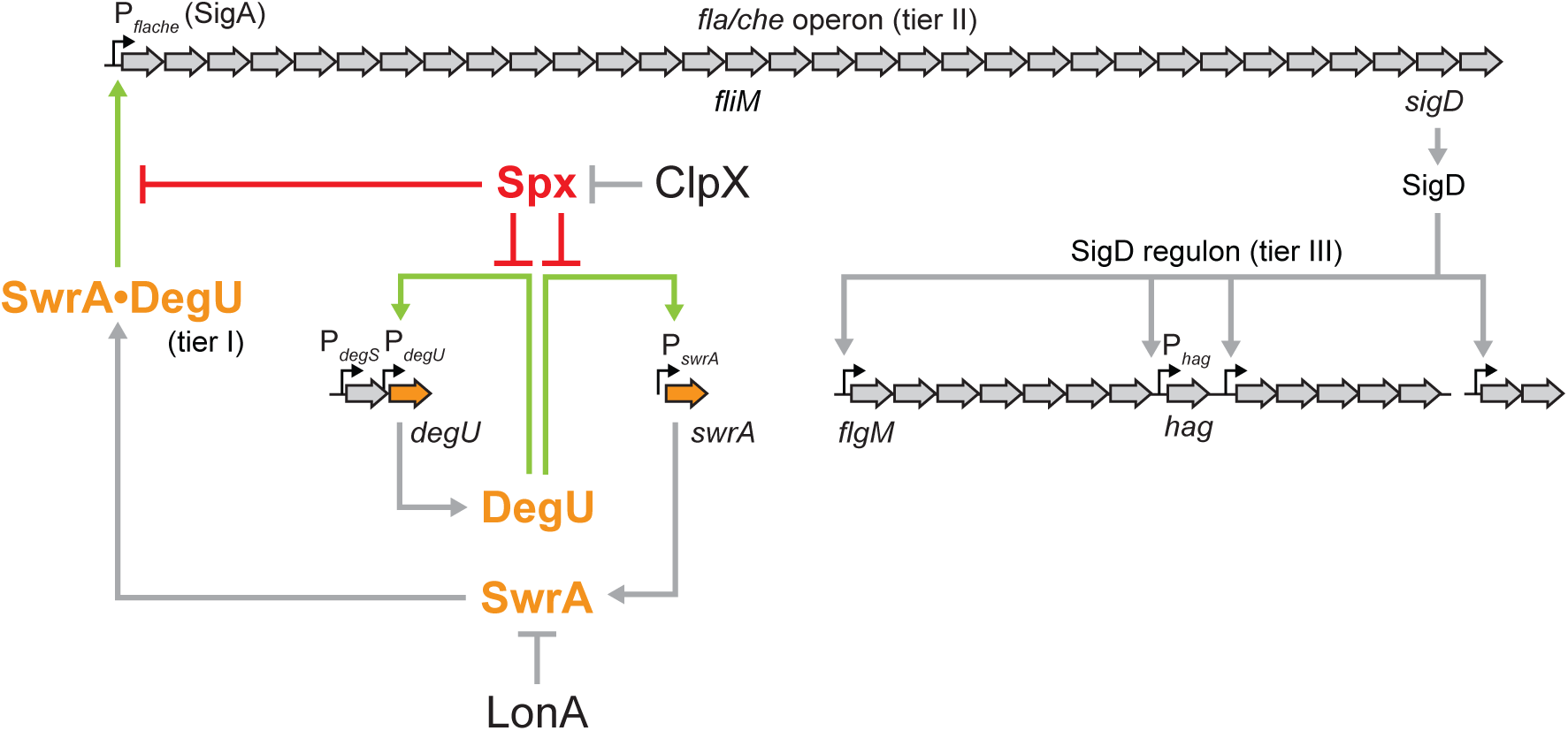
A cartoon diagram of the flagellar regulon of *Bacillus subtilis*. The flagellar regulon of *B. subtilis* is organized in three tiers. Tier I is the SwrA•DegU hybrid master activator that activates expression of the sigma A dependent P*_fla/che_* promoter. Tier II is the *fla/che* operon that contains thirty-two genes dedicated to the assembly of the flagellar basal body, rod and hook, as well as genes for chemotaxis signal transduction and the alternative sigma factor SigD. Tier III is the SigD regulon (shown abbreviated) containing proteins involved in flagellar filament assembly and flagellar rotation (among other products). Block arrows indicate genes and the names of genes mentioned in text are indicated in italics below their corresponding coding region. Small bent arrows indicate promoters and the names of the promoters mentioned in text are indicated above the promoter location. Thick arrows indicate activation, T bars indicate inhibition. Light green arrows indicate DegU-dependent transcriptional activation; red T-bars indicate Spx-dependent inhibition.

One mutation that enhances swarming motility is a mutation of the gene *lonA*, which encodes the cytoplasmic protease LonA (Chen 2009). LonA degrades SwrA during growth in liquid media and requires the adaptor protein SmiA, which both primes SwrA for degradation and delivers it to the LonA protease (Mukherjee 2015; Hughes 2018; Olney 2022). When cells are inoculated on a surface conducive to swarming, SwrA levels increase seemingly due to the inactivation of proteolysis (Mukherjee 2015). In turn, a complex of SwrA•DegU binds to and activates the promoter of the *fla/che* operon, flagellar biosynthesis is activated, flagellar density on the cell surface increases, and swarming commences (Kearns 2005; Ogura 2012, Mordini 2013; Guttenplan 2013; Mishra 2024). How exactly surface contact antagonizes SwrA proteolysis is unknown but the inhibitory role of swarming motility is specific to LonA, as mutation of other cytoplasmic proteolysis systems in the cell do not similarly enhance swarming motility (Mukherjee 2015). Indeed ClpX, one of several unfoldase partners for the cytoplasmic protease ClpP (Gottesman 1993; Kim 2000), promotes motility such that both swimming and swarming are abolished when it is mutated (Mukherjee 2015; Moliere 2016).

Previous work indicated that mutation of ClpX abolished swimming motility in laboratory strains indirectly through the global transcription factor Spx (Moliere 2016). Spx is restricted to very low levels in the cell by ClpX-mediated proteolysis but accumulates to a high level when ClpX is mutated (Nakano 2001; Nakano 2002). Spx is an unusual transcription factor in that it acts as a monomer to both activate and repress gene expression (Lin 2012; Nakano and Kuester 2003). Spx activates transcription by interacting with the alpha subunit of RNA polymerase and the vegetative sigma factor SigA, and is thought to enhance binding of the RNAP holoenzyme to certain promoters based on a short sequence of DNA upstream of the -35 element (Reyes 2008; Nakano 2005; Nakano 2010; Lin 2013; Shi 2021). Spx is also thought to inhibit transcription by binding to the alpha subunit and neutralizing interaction with other transcriptional activators, but the need for a DNA binding sequence in this case is unclear (Nakano 2003; Newberry 2005). Spx controls a wide variety of processes in *B. subtilis*, including activation of the oxidative and/or disulphide stress response pathway (Gaballa 2013, Leelakriansgsak 2007; Choi 2006; Nakano and Erwin 2005), activation of cell division (Yu 2021), inhibition of surfactin gene expression (Nakano 2003, Nakano and Kuester 2003), inhibition of sulfur uptake (Erwin 2005), and the inhibition of competence for DNA import (Nakano 2001; Rochat 2012). Relatively few promoters that are directly antagonized by Spx have been identified, and the target responsible for motility inhibition is unknown.

Here we show that mutation of ClpX abolishes swarming motility in *B. subtilis* due to a defect in flagellar number. We track the motility defect to a decrease in the level of transcription of the *fla/che* operon, a result consistent with previous reports in laboratory strains (Moliere 2016). Further we find that in the absence of ClpX, protein levels of the master motility regulators DegU and SwrA are reduced, suggesting that the defect occurs upstream of *fla/che* operon transcription. Suppressors that restored swarming to cells mutated for *clpX* had additional mutations in either *lonA* or *spx*. While both classes of suppressors increased the expression of the *degU* and *swrA* genes, we infer that LonA primarily acts by restricting SwrA protein accumulation and that Spx primarily acts by reducing transcription of DegU-activated genes. Spx levels rise under cell envelope stress and we suggest that Spx-mediated repression of motility may serve to limit stress associated with flagellar biosynthesis.

## MATERIALS AND METHODS

### Growth conditions

*B. subtilis* strains were grown in lysogeny (LB) broth (10 g tryptone, 5 g yeast extract, 5 g NaCl per L) or on LB plates fortified with 1.5% Bacto agar at 37°C. When appropriate, antibiotics were included at the following concentrations: 5 µg/ml chloramphenicol (*cat*), 5 µg/ml kanamycin (*kan*), 100 µg/ml spectinomycin (*spec*), 10 µg/ml tetracycline (*tet*), or 1 µg/ml erythromycin plus 25 µg/ml lincomycin (*mls*).

### SPP1 phage transduction

To 200 µL of dense culture grown in TY broth (LB broth supplemented after autoclaving with 10 mM MgSO_4_ and 100 µM MnSO_4_), serial dilutions of SPP1 phage stock were added. To each mixture, 3 ml molten TY top agar (TY supplemented with 0.5% agar) was added, the mixture was poured atop fresh TY plates (TY fortified with 1.5% agar), and the plates were incubated at 30°C overnight. Top agar from a plate containing near confluent plaques was harvested by scraping into a 15 ml conical tube with 5 ml of TY broth, vortexed, and centrifuged at 6,500 x g for 5 min. The supernatant was passed through a 0.2 µm syringe filter and stored at 4°C.

Recipient cells for transduction were grown to stationary phase in 3 ml TY broth at 37°C. 1 ml of culture was mixed with 25 µl of SPP1 donor phage stock made above. 9 ml of TY broth was added to the mixture and allowed to stand at 37°C for 20 min. The transduction mixture was then centrifuged at 6,000 x g for 5 min, the supernatant was discarded, and the pellet was resuspended in the remaining volume. Cell suspension (∼100 µl) was then plated on LB plates fortified with the appropriate antibiotic for selection and 10 mM sodium citrate if the antibiotic was chloramphenicol, kanamycin, or spectinomycin.

### Strain construction

All constructs were either first introduced into the domesticated *Bacillus subtilis* strain PY79 by natural competence and then transferred to the undomesticated *Bacillus subtilis* 3610 background using SPP1-mediated generalized phage transduction (Yasbin and Young, 1974) or transformed directly into DK1042 (Konkol 2013). All plasmids used in this study are listed in Supplemental Table S1. All primers used in this study are listed in Supplemental Table S2. All strains used in this study are listed in Table 2.

**TABLE 1:**
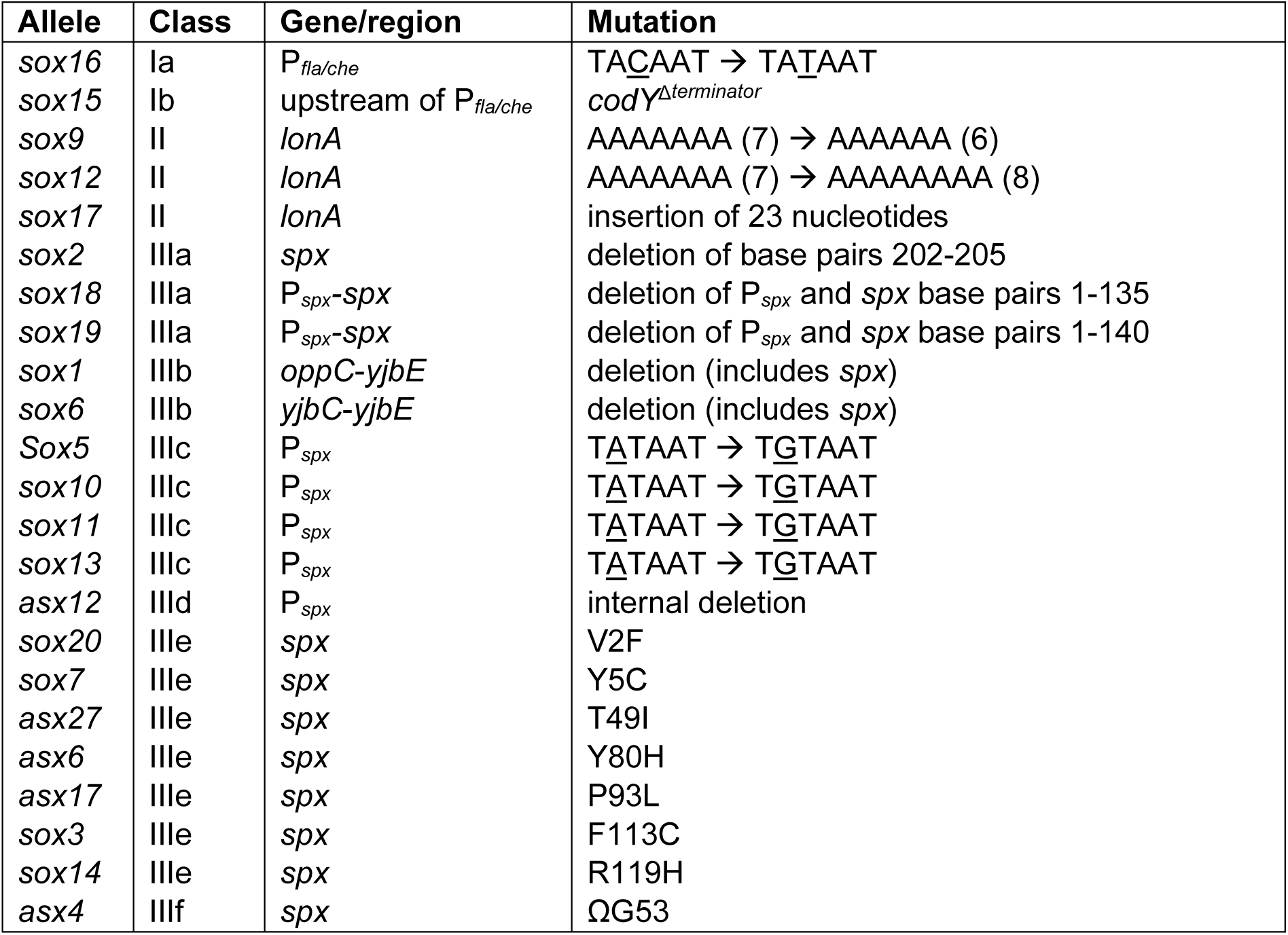
Suppressor mutations that restored swarming in the absence of ClpX.

**TABLE 2:**
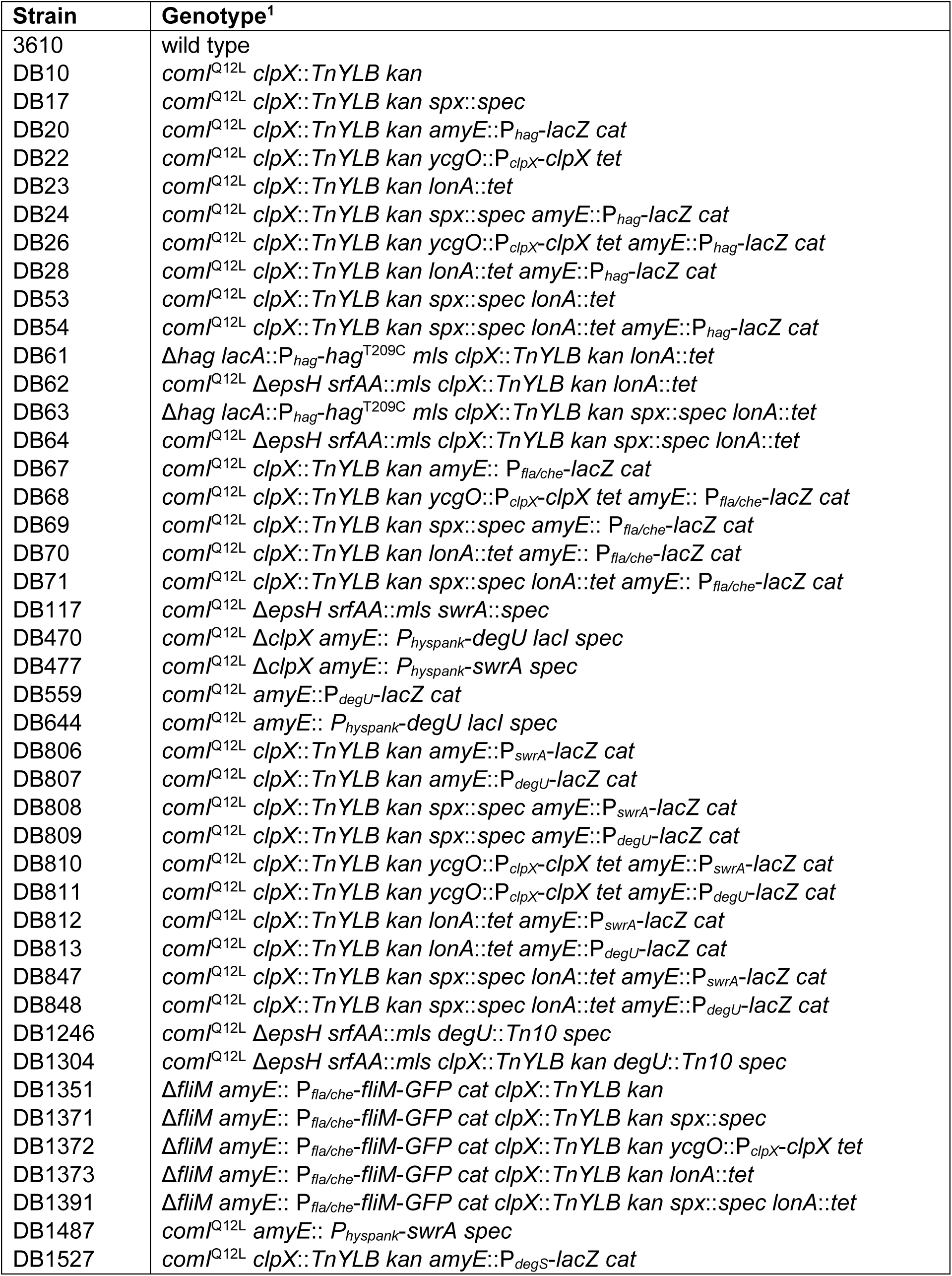
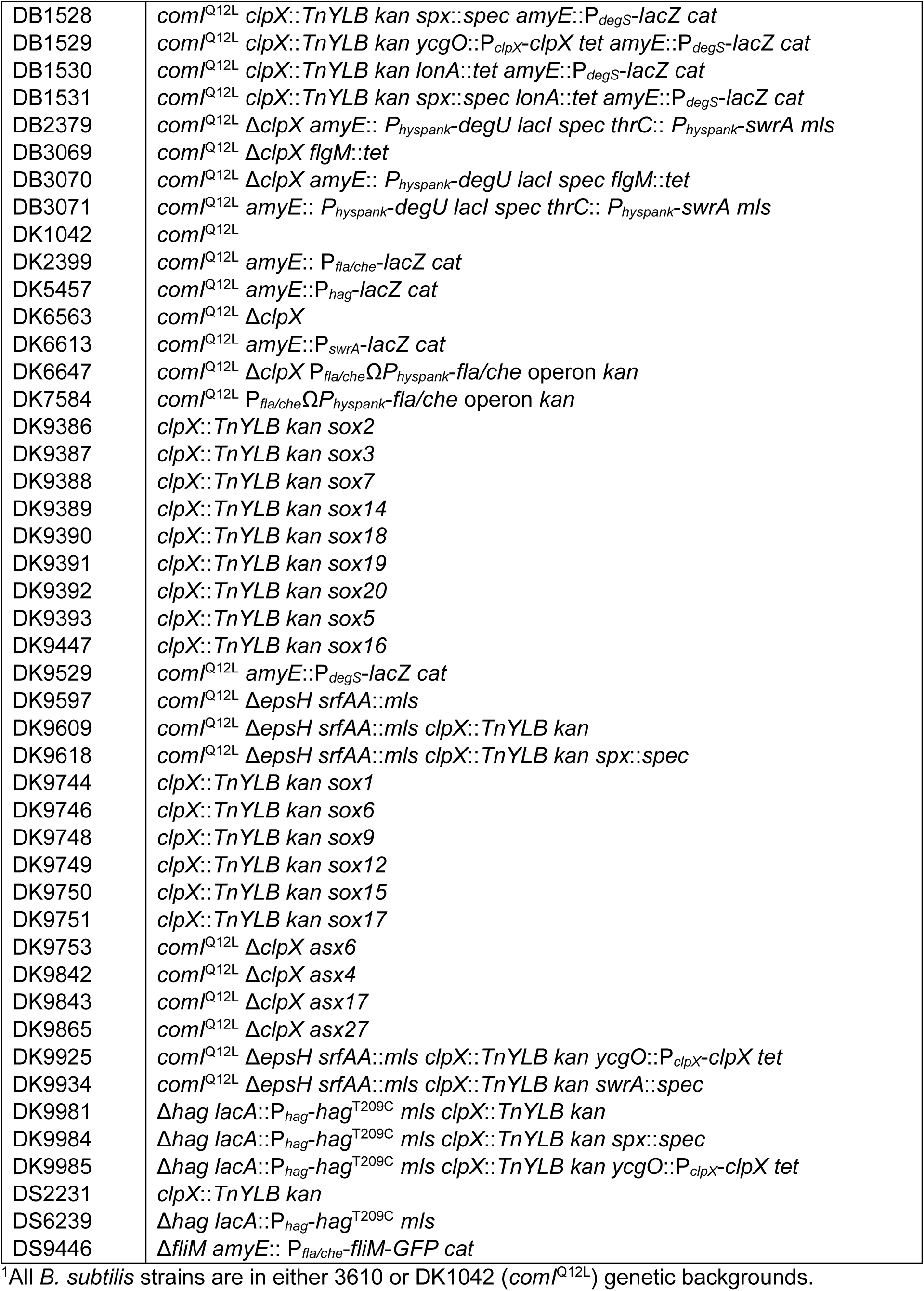
Strains.

#### Antibiotic replacement constructs

The *spx*::*spec* allele was a generous gift from John Helmann (Cornell University).

The *lonA::tet* insertion deletion allele was generated using a modified “Gibson” isothermal assembly protocol (Gibson et al. 2009). Briefly, the region upstream of *lonA* was PCR amplified using the primer pair 7838/7839 and the region downstream of *lonA* was PCR amplified using the primer pair 7840/7841. Primers 7839 and 7840 contained a short sequence (23 bp) of homology to the tetracycline resistance gene. DNA containing a tetracycline resistance gene was amplified using universal primers 3250/3251 and pDG1515 as a template (Guérout-Fleury 1995). The three DNA fragments were combined at equimolar amounts to a total volume of 5 µL and added to a 10 µl aliquot of 2X prepared master mix (see below). The reaction was incubated for 60 minutes at 50°C. The completed reaction was then PCR amplified using primers 7838/7841 to amplify the assembled product. The amplified product was transformed into competent cells of PY79 and then transferred to the 3610 background using SPP1-mediated generalized transduction and selection on tetracycline. The 5X isothermal assembly reaction buffer (500 mM Tris-HCl pH 7.5, 50 mM MgCl_2_, 1 mM of each dNTP (New England BioLabs), 50 mM DTT (Bio-Rad), 312.5 mM PEG-8000 (Fisher Scientific), and 5 mM NAD^+^ (New England BioLabs)) was aliquoted and stored at -80°C. The 2X assembly master mixture was made by combining the prepared 5X isothermal assembly reaction buffer with T5 exonuclease diluted 1:5 with 5X reaction buffer (New England BioLabs) (0.01 units/µL), Phusion DNA polymerase (New England BioLabs) (0.033 units/µL), and Taq DNA ligase (New England BioLabs) (5328 units/µL). The master mix was aliquoted as 10 µl and stored at -80°C.

#### In-frame deletions

To generate the Δ*clpX* in-frame markerless deletion construct, the region upstream of *clpX* was PCR amplified using the primer pair 6333/6334 and 6335/6336 using 3610 chromosomal DNA as a template. The PCR products was then assembled by isothermal assembly, the resulting assembly was digested with BamHI/KpnI, and cloned into the BamHI/KpnI sites of pMiniMAD2 which carries a temperature sensitive origin of replication and a *mls* resistance cassette (Patrick 2008) to generate pSO18. The plasmid pSO18 was purified from *recA*^+^ TG1 *E. coli* and introduced to DK1042 by transformation at the restrictive temperature for plasmid replication (37°C) using *mls* resistance as a selection. To evict the plasmid, the strain was incubated in 3 ml LB broth at a permissive temperature for plasmid replication (22°C) for 14 hours, then serially diluted and plated on LB agar at 37°C. Individual colonies were patched on LB plates and LB plates containing *mls* to identify *mls*-sensitive colonies that had evicted the plasmid (Arnaud 2004). Chromosomal DNA from colonies that had excised the plasmid was purified and screened by PCR using primers 6333/6336 to determine which isolate had retained the Δ*clpX* allele.

#### Complementation constructs

To generate the P*_clpX_-clpX* complementation construct pDP538, a PCR product containing the *clpX* coding region plus 434 base pairs of upstream sequence was amplified from *B. subtilis* 3610 chromosomal DNA using the primer pair 7410/7411, digested with *Xho*I and *Nhe*I and cloned into the *Xho*I and *Nhe*I sites of pKM86 containing a polylinker and tetracycline resistance cassette between two arms of *ycgO* (generous gift of Dr. David Rudner, Harvard Medical School).

#### Inducible constructs

The *P_hyspank_-degU* construct was the generous gift of Dr. Nicola Stanley-Wall (University of Dundee).

#### Transcriptional reporter constructs

To generate the β-galactosidase (*lacZ*) reporter constructs pAEJ5, pAEJ9, and pAEJ15, PCR products containing the following promoters were amplified from *B. subtilis* 3610 or DK1042 chromosomal DNA using the primers indicated in parentheses: P*_degS_* (7657/7658) and P*_degU_* (7944/7945). Each PCR product was digested with *Eco*RI and either *Bam*HI or *Hind*III and cloned independently into the *Eco*RI and either *Bam*HI or *Hind*III sites of plasmid pDG268, which carries a chloramphenicol resistance marker and a polylinker upstream of the *lacZ* gene between two arms of the *amyE* gene (Antoniewski 1990).

### Swarm expansion assay

Cells were grown overnight at room temperature in LB broth, back-diluted and grown to mid-log phase at 37°C in LB broth and 1mM IPTG (if applicable), and resuspended to an OD_600_ of 10 in MQ H_2_O containing 0.5% India ink. Freshly prepared LB plates containing 0.7% Bacto agar (25 ml/plate) and 1mM IPTG (if applicable) were dried for 10 minutes in a laminar flow hood, centrally inoculated with 10 µl of the cell suspension, dried for another 10 minutes, and incubated at 37°C for 6 hours. Each strain was done in technical triplicate. The India ink demarcates the origin of the inoculation, and the swarm radius was measured in mm relative to this origin. For consistency, an axis was drawn on the back of the plate and swarm radii measurements were taken along this axis.

### Swim assay

Swim agar plates containing 25 ml LB fortified with 0.3% Bacto agar were prepared fresh. Each swim plate was toothpick-inoculated from a colony grown overnight on a plate and photographed for motility after 12 hours of incubation at 37°C. Plates were visualized with a camera.

### Microscopy

Fluorescence microscopy was performed with a Nikon 80i microscope with a phase contrast objective Nikon Plan Apo 100X and an Excite 120 metal halide lamp. FM4-64 was visualized with a C-FL HYQ Texas Red Filter Cube (excitation filter 532-587 nm, barrier filter >590 nm). GFP or fluorescent maleimide stain was visualized using a C-FL HYQ FITC Filter Cube (excitation filter 460-500 nm, barrier filter 515-550 nm). Images were captured with a Photometrics Coolsnap HQ2 camera in black and white using NIS-Elements software and subsequently false-colored and superimposed using Fiji v2.1.0 (Schindelin, 2012).

For microscopy of strains expressing GFP, cells were grown in LB broth at 37°C to an OD_600_ of 0.6-0.85. 1 ml of culture was pelleted, and the pellet was resuspended in 30 µl of 1X phosphate-buffered saline (PBS) containing 5 µg/ml FM 4-64 (Invitrogen #T13320) and incubated for 2 min in the dark at room temperature. Cells were washed with 1 ml 1X PBS and pelleted again. Cells were then resuspended in 30 µl 1X PBS. Cells were observed by spotting 4 µl of the resuspension on a glass microscope slide and immobilized with a poly-L-lysine-treated glass coverslip. Images were captured with NIS-Elements software.

For fluorescent microscopy of flagella, cells were grown in LB broth at 37°C to an OD_600_ of 0.6-0.85. 1 ml of culture was pelleted, and the pellet was resuspended in 50 µl of 1X PBS containing 5 µg/ml Alexa Fluor 488 C_5_ maleimide (Invitrogen; molecular probes A10254) and incubated for 3 min in the dark at room temperature. Cells were washed with 1 ml 1X PBS and pelleted again. The pellet was resuspended in 30 µl of 1X phosphate-buffered saline (PBS) containing 5 µg/ml FM 4-64 and incubated for 2 min in the dark at room temperature. Cells were washed with 1 ml 1X PBS and pelleted again. Cells were then resuspended in 30 µl 1X PBS. Cells were observed by spotting 4 µl of the resuspension on a glass microscope slide and immobilized with a poly-L-lysine-treated glass coverslip. Images were captured with NIS-Elements software.

### Western blotting

*B. subtilis* strains were grown in LB to an OD_600_ of ∼1 either from colony inoculation or back-dilution from overnight cultures. 1 ml (or (10 ml) was harvested by centrifugation, and pellets were resuspended to an OD_600_ of 10 (or OD_600_ of 100 if 10 ml samples) in lysis buffer (50 mM Tris pH 7.0, 100 µg/ml lysozyme, 10 µg/ml DNAse I, 100 µg/ml RNAse A, 1 mM phenylmethylsulfonyl fluoride (PMSF), 10 mM MgCl_2_) and incubated at 37°C for 1 hour. 6x sodium dodecyl sulfate (SDS) loading dye supplemented with β-mercaptoethanol (βME) was added to a 1x concentration. Samples were separated by 12 or 15% SDS-polyacrylamide gel electrophoresis (SDS-PAGE). The proteins were electroblotted onto nitrocellulose and the membrane was blocked with 5% skim milk in tris-buffered saline with Tween 20 (TBST) and developed with varying dilutions of primary antibody (1:80,000 for α-Hag, 1:10,000 for α-SigD, 1:10,000 for α-DegU, 1:2,000 for α-SwrA, 1:40,000 for α-SigA) and a 1:10,000 dilution secondary antibody (horseradish peroxidase-conjugated goat anti-rabbit immunoglobulin G). 10 ml samples were used to blot for SigD and DegU, while 1 ml samples were used for the rest. Immunoblot was developed using the Immun-Star HRP developer kit (Bio-Rad).

### β-galactosidase assays

Three biological replicates of each strain were grown in 3 ml of LB broth at 37°C to an OD_600_ of 0.7-1.0. 200 µl of culture was pelleted, and the pellet was resuspended in 200 µl of Z-buffer (16.1 g Na_2_HPO_4_ • 7H_2_O + 5.5 g NaH_2_PO_4_ • H_2_O + 0.75 g KCl + 0.246 g MgSO_4_ • 7H_2_O in 1 L H_2_O, pH = 7) with 0.27% BME. 4 µl of 10 mg/ml lysozyme was added, and samples were incubated at 30°C for 15 min to lyse. In a 96-well plate, lysates were diluted in Z-buffer to 200 µl depending on the strength of the promoter fused to *lacZ* (P*_hag_* = 1:25, P*_fla/che_* = 1:10, P*_degS_* = 1:2, P*_degU_* = 1:1 (undiluted), P*_swrA_* = 1:2). Three wells had 200 µL of Z-buffer only and served as negative controls. 40 µl of 4 mg/ml ortho-nitrophenyl-β-D-galactopyranoside (ONPG) in Z-buffer was added to each reaction. The plate was incubated at 30°C for 1 hr, and the OD_420_ and OD_550_ of each well were taken every 2 min. The slope of each sample’s ODs over time were derived. The average slope of the three negative control wells was subtracted from the slope of each experimental well. The Miller Units were calculated using the following formula:

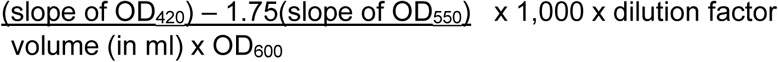

### Sequencing candidate genes

PCR product containing Pfla/che was amplified from chromosomal DNA from DS2231 suppressor *sox16* using the primers 1921/3042. The P*_fla/che_* PCR product was then sequenced using primers 1921 and 3042 individually. PCR product containing P*_spx_* was amplified from DS2231 suppressors *sox5, 10-11,* and *13* and from DK6563 suppressor *asx12* using the primers 6589/6590. The P*_spx_* PCR product was then sequenced using primers 6589/6590. PCR product containing *spx* was amplified from DS2231 suppressors *sox2-3, 7, 14,* and *18-20* and from DK6563 suppressors *asx4, 6, 16,* and *27* using the primers 6589/6590. The *spx* PCR product was then sequenced using primers 6591/6590.

### Whole-genome sequencing

One colony of each strain was grown overnight in 3 ml of LB broth at room temperature. 2 ml of culture per strain was harvested, and the pellets were each resuspended in 180 µl enzymatic lysis buffer (ELB; 20 mM Tris-HCl pH 8, 2mM ethylenediaminetetraacetic acid (EDTA), 1.2% Triton X-100). The samples were incubated at 37°C for 30 min. 10 µl of 4 mg/ml RNAse A was added and mixed by pipetting. The samples were incubated at 37°C for 30 min. 25 µl of Proteinase K was added and mixed by vortexing for 30 sec. The samples were incubated at 37°C for 10 min. 200 µl of Buffer AL was added and mixed by vortexing for 30 sec. The samples were incubated at 56°C for 30 min. 200 µl of 100% ethanol was added and mixed by vortexing for 30 sec. The entire sample was pipetted onto a DNeasy mini spin column in a 2 ml collection tube and centrifuged at 10,000 x g for 1 min. The collection tube was discarded and placed into a fresh one. 500 µl of Buffer AW1 was added to the column, and the column was centrifuged at 10,000 x g for 1 min. The collection tube was discarded and placed into a fresh one. 500 µl of Buffer AW2 was added to the column, and the column was centrifuged at maximum speed for 3 min. The flow-through was discarded, and the column was replaced and centrifuged again at maximum speed for 1 min to remove residual ethanol. The collection tube was discarded and placed into a fresh Eppendorf tube. 100 µl of Buffer AE was added to the column, and the column was incubated at room temperature for 2 min. The column was centrifuged at 10,000 x g for 1 min to elute DNA. The elution step was repeated to elute additional DNA, for a total of 200 µl. An additional RNAseA treatment was performed by adding 4 µl of 4 mg/ml RNAse A and incubating at room temperature for 10 min. Samples were submitted to Center for Genomics and Bioinformatics at Indiana University for library preparation and data analysis.

## RESULTS

### ClpX promotes swarming motility by enhancing flagellar biosynthesis

The ClpX unfoldase subunit of the ClpP protease has been shown to be required for swimming motility through soft agar plates in *B. subtilis* laboratory strains (Moliere 2016). Consistent with a role in promoting flagellar motility, mutation of the *clpX* gene abolished swarming (**Fig 2A**) (Sampriti 2015; Sanchez 2022) and reduced swimming in the NCIB3610 ancestral strain background (**Fig 3**), and both phenotypes were restored by complementation when *clpX* was expressed from its native promoter and inserted at an ectopic site in the chromosome (**Fig 2A**, **Fig 3**). We conclude that ClpX is required for swarming and enhances swimming motility. We note that mutation of *clpX* in the ancestral strain did not completely abolish swimming through a soft agar matrix as previously reported, and we infer that the difference in phenotype may be due to genetic differences between strain backgrounds.

**Figure 2:**
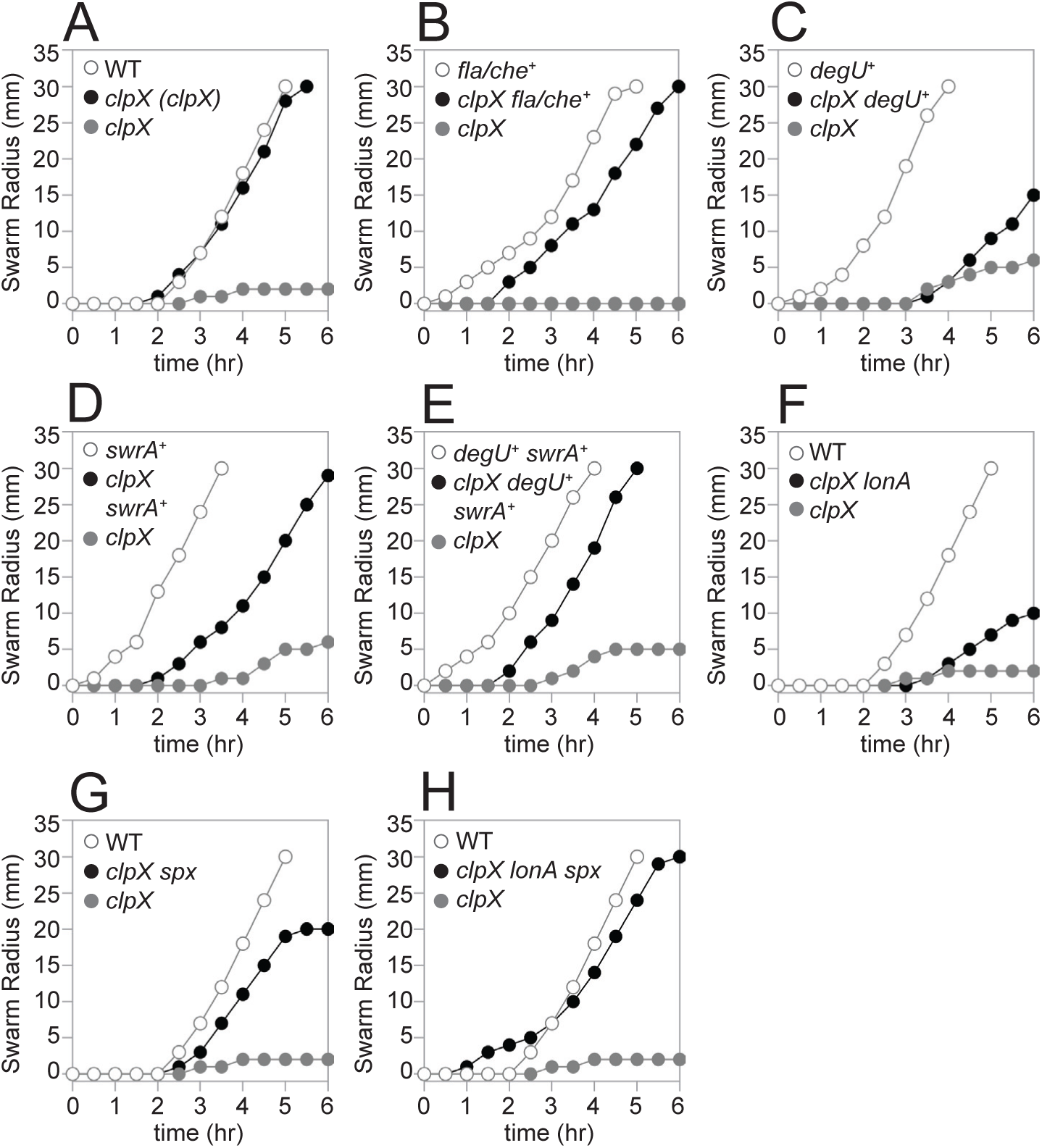
Mutation of LonA and Spx restores swarming in the absence of ClpX. Quantitative swarm expansion assays for the strains indicated in each panel. Genes in italics are mutated, genes in parenthesis are complementation constructs expressed under the gene’s native promoter and inserted at an ectopic site in the chromosome, and a “+” indicates that the gene was induced with 1 mM IPTG. Each data point is the average of three technical replicates. (A) The *clpX* mutant phenotype was complemented by reintroducing *clpX* under its native promoter at an ectopic site. The following strains were used to generate the data in this panel: WT (DK1042), *clpX (clpX)* (DB22), *clpX* (DB10). (B) The *clpX* mutant phenotype was rescued when the *fla/che* operon was artifically overexpressed. The following strains were used to generate the data in this panel: *fla/che*^+^ (DK7584, 1 mM IPTG added), *clpX fla/che*^+^ (DK6647, 1 mM IPTG added), *clpX* (DK6647, 0 mM IPTG added). (C) The *clpX* mutant phenotype was partially rescued when the *degU* gene was artifically overexpressed. The following strains were used to generate the data in this panel: *degU*^+^ (DB644, 1 mM IPTG added), *clpX degU*^+^ (DB470, 1 mM IPTG added), *clpX* (DB470, 0 mM IPTG added). (D) The *clpX* mutant phenotype was partially rescued when the *swrA* gene was artifically overexpressed. The following strains were used to generate the data in this panel: *swrA*^+^ (DB1487, 1 mM IPTG added), *clpX swrA*^+^ (DB477, 1 mM IPTG added), *clpX* (DB477, 0 mM IPTG added). (E) The *clpX* mutant phenotype was rescued when both the *degU* and *swrA* genes were artifically overexpressed. The following strains were used to generate the data in this panel: *degU*^+^ *swrA*^+^ (DB3071, 1 mM IPTG added), *clpX degU*^+^ *swrA*^+^ (DB2379, 1 mM IPTG added), *clpX* (DB2379, 0 mM IPTG added). (F) The *clpX* mutant phenotype was partially rescued when the *lonA* gene was also mutated. The following strains were used to generate the data in this panel: WT (DK1042), *clpX lonA* (DB23), *clpX* (DB10). (G) The *clpX* mutant phenotype was partially rescued when the *spx* gene was also mutated. The following strains were used to generate the data in this panel: WT (DK1042), *clpX spx* (DB17), *clpX* (DB10). (H) The *clpX* mutant phenotype was fully rescued when both the *lonA* and *spx* genes were also mutated. The following strains were used to generate the data in this panel: WT (DK1042), *clpX lonA spx* (DB53), *clpX* (DB10).

**Figure 3:**
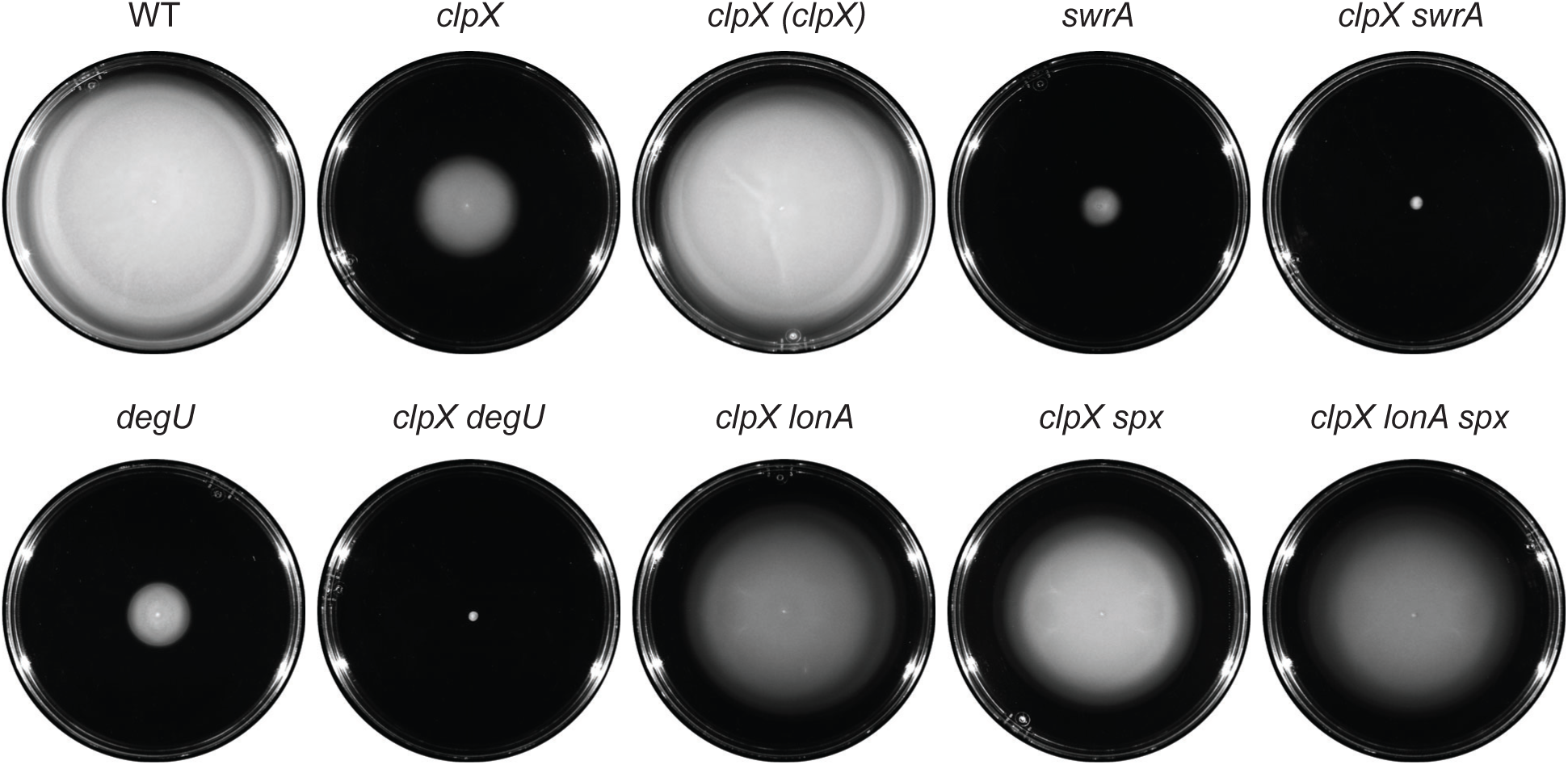
Mutation of LonA and Spx restores swimming in the absence of ClpX. Cells were centrally inoculated by toothpick on 0.3% agar swim plates, incubated for 12 hours at 37°C, and filmed against a black background such that uncolonized agar appears black and bacterial growth appears white. Genes in italics are mutated, and genes in parentheses are complementation constructs expressed under the gene’s native promoter and inserted at an ectopic site in the chromosome. Each strain is additionally mutated for *srfAA* and *epsH* to abolish the contribution of sliding motility on the surface. The following strains were used to generate the data in this figure: WT (DK9597), *clpX* (DK9609), *clpX* (*clpX)* (DK9925), *swrA* (DB117), *clpX swrA* (DK9934), *degU* (DB1246), *clpX degU* (DB1304), *clpX lonA* (DB62), *clpX spx* (DK9618), *clpX lonA spx* (DB64).

One potentially relevant genetic locus that could explain the differential consequence of a *clpX* mutation in ancestral and laboratory strains is the gene *swrA*, which encodes the master flagellar activator component SwrA. Laboratory strains are naturally defective for SwrA, whereas ancestral strains are not (Kearns 2004; Calvio 2005, Zeigler 2008). Mutation of *swrA* reduced swimming in the ancestral background (**Fig 3**) and cells doubly mutated for both *swrA* and *clpX* recapitulated the more severe *clpX* mutant defect in laboratory strain swimming (**Fig 3**) (Moliere 2016). SwrA cooperates with the response regulator DegU to activate flagellar gene expression, and like mutation of SwrA, mutation of *degU* alone reduced swimming and further exacerbated the *clpX* mutant phenotype when simultaneously mutated (**Fig 3**). We conclude that ClpX enhances flagellar motility in *B. subtilis,* the extent of which depends on the genetic background.

One way in which a *clpX* mutant could be defective in swarming is if it failed to produce surfactin. Indeed, mutation of *clpX* has been shown to result in the inhibition of the promoter for the *srf* operon responsible for surfactin biosynthesis (Nakano 1991; Nakano 2000). On swarm agar plates however, a large zone of clear watery surface indicative of surfactin production was observed to surround the non-swarming colony (Kearns 2003; Dubois 2008). It is unclear why cells mutated for ClpX appeared to produce abundant surfactin, but we note that previous work documenting reduced surfactin gene expression was performed in domesticated laboratory strains that lack surfactin production, and the presence of the ancestral allele of *sfp* alters surfactin gene expression (Nakano 1992). Whatever the case with regards to *srf* gene expression, we conclude that swarming is likely not limited by surfactin production in the absence of ClpX.

Another reason that the *clpX* mutant might be impaired for swarming motility is if it assembled fewer flagella on the cell surface (Mukherjee 2015; Phillips 2015). To determine whether flagellar number was reduced in the absence of ClpX, a version of the flagellar filament protein Hag which could be fluorescently labeled by the addition of a maleimide stain (Hag^T209C^) was introduced (Blair 2008). Mutation of *clpX* appeared to dramatically reduce flagellar density per cell relative to wild type (**Fig 4A**). Moreover, mutation of *clpX* decreased Hag protein accumulation in Western blot analysis (**Fig 5**) and also reduced expression of a transcriptional reporter in which the *hag* promoter, P*_hag_*, was fused to the *lacZ* gene encoding β-galactosidase (**Fig 6A**). For each assay, wild type phenotypes were restored when *clpX* was ectopically complemented (**Fig 4A**, **Fig 5**, **Fig 6A**). We conclude that the swarming defect in the absence of ClpX is correlated with reduced transcription from the P*_hag_* promoter which, in turn, reduces the filament protein pool and restricts flagellar filament assembly.

**Figure 4:**
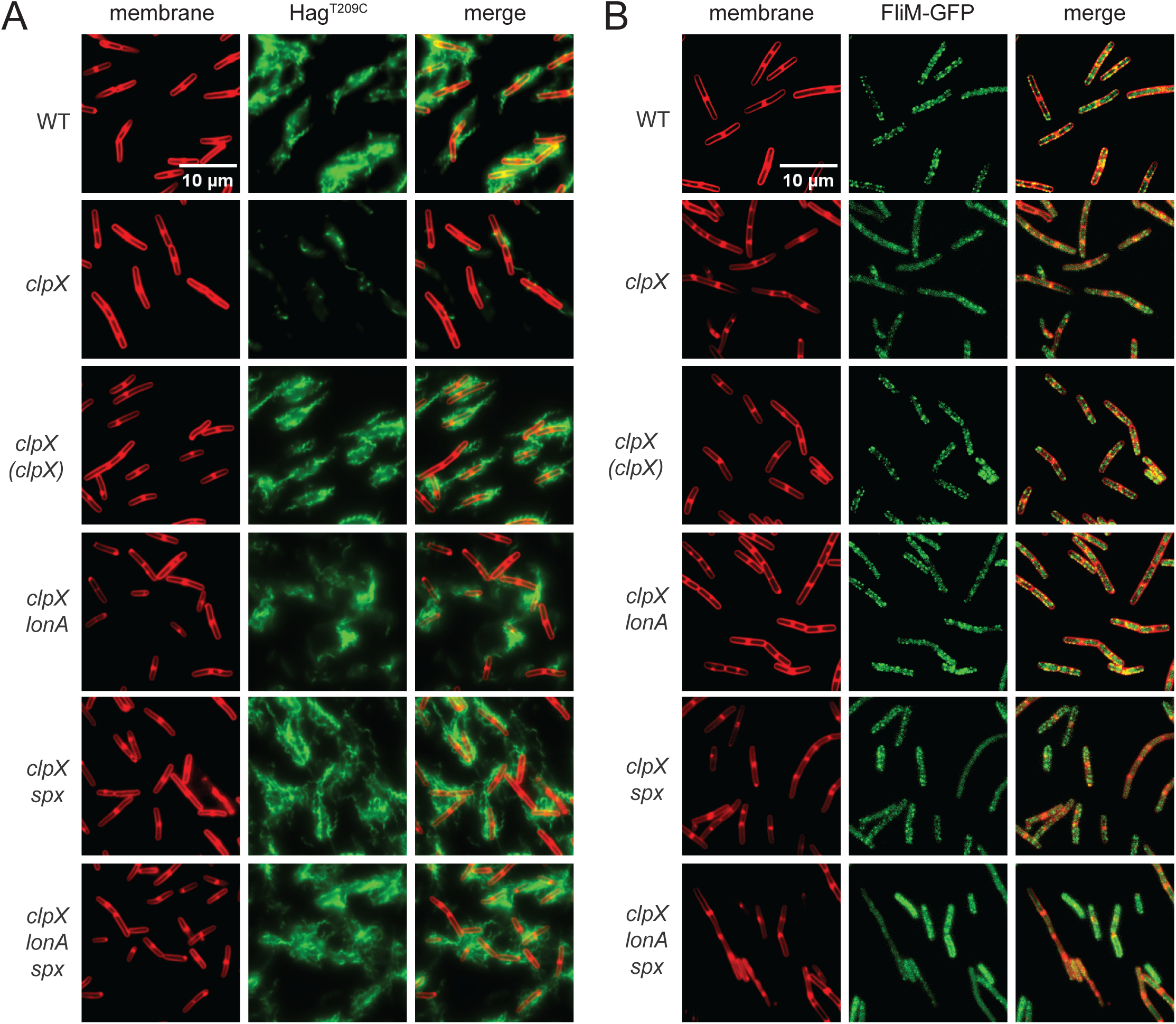
Cells mutated for ClpX have a reduced number of flagella per cell. Fluorescence microscopy observation of the cell membrane and flagella (A) or basal bodies (B) in a variety of genetic backgrounds. Genes in italics are mutated, and genes in parentheses are complementation constructs expressed under the gene’s native promoter and inserted at an ectopic site in the chromosome. (A) Fluorescent FM 4-64 membrane (false-colored red) and Alexa Fluor maleimide of Hag^T209C^ (false-colored green) staining in a variety of genetic backgrounds. Each strain is additionally mutated for the native copy of the *hag* gene and has the *hag*^T209C^ allele expressed under the native promoter at an ectopic site in the chromosome. The following strains were used to generate the data in this panel: WT (DS6239), *clpX* (DK9981), *clpX* (*clpX)* (DK9985), *clpX lonA* (DB61), *clpX spx* (DK9984), *clpX lonA spx* (DB63). (B) Fluorescent FM 4-64 membrane staining (false-colored red) and fluorescent FliM-GFP translational fusion (false-colored green) in a variety of genetic backgrounds. Each strain is additionally mutated for the native copy of the *fliM* gene and has the *fliM*-*GFP* allele expressed under the native promoter at an ectopic site in the chromosome. The following strains were used to generate the data in this panel: WT (DS9446), *clpX* (DB1351), *clpX* (*clpX)* (DB1372), *clpX lonA* (DB1373), *clpX spx* (DB1371), *clpX lonA spx* (DB1391).

**Figure 5:**
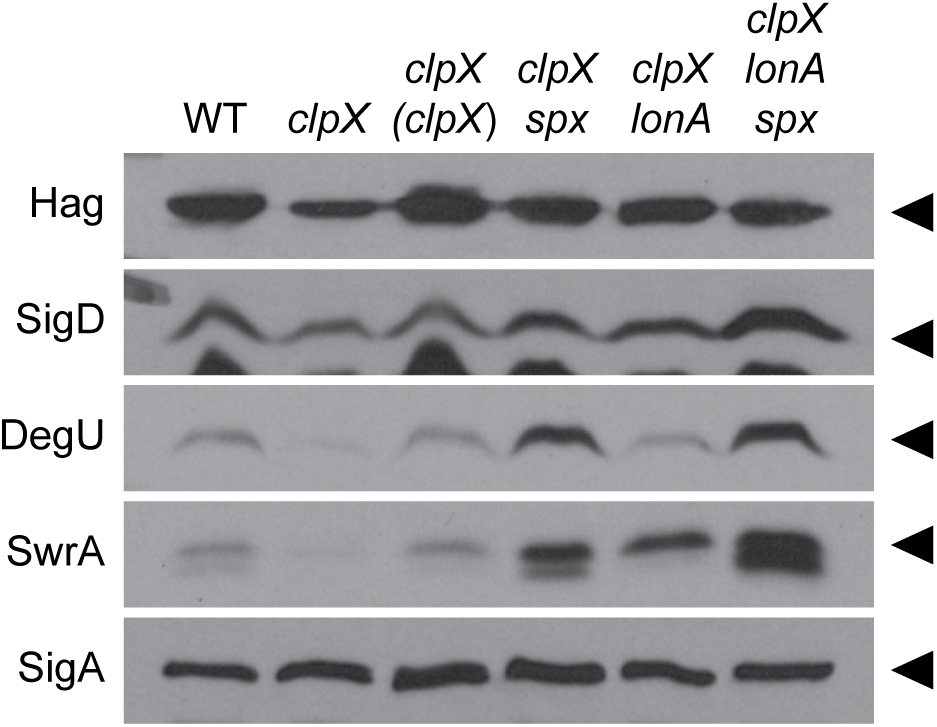
Mutation of LonA and Spx increases levels of key motility proteins in the absence of ClpX. Western blot analysis with the primary antibody indicated on the left and the genetic background indicated above. Genes in italics are mutated, and genes in parentheses are complementation constructs expressed under the gene’s native promoter and inserted at an ectopic site in the chromosome. Carets on the right indicate the location of the corresponding protein. The following strains were used to generate the data in this figure: WT (DK1042), *clpX* (DB10), *clpX* (*clpX)* (DB22), *clpX spx* (DB17), *clpX lonA* (DB23), *clpX lonA spx* (DB53).

**Figure 6:**
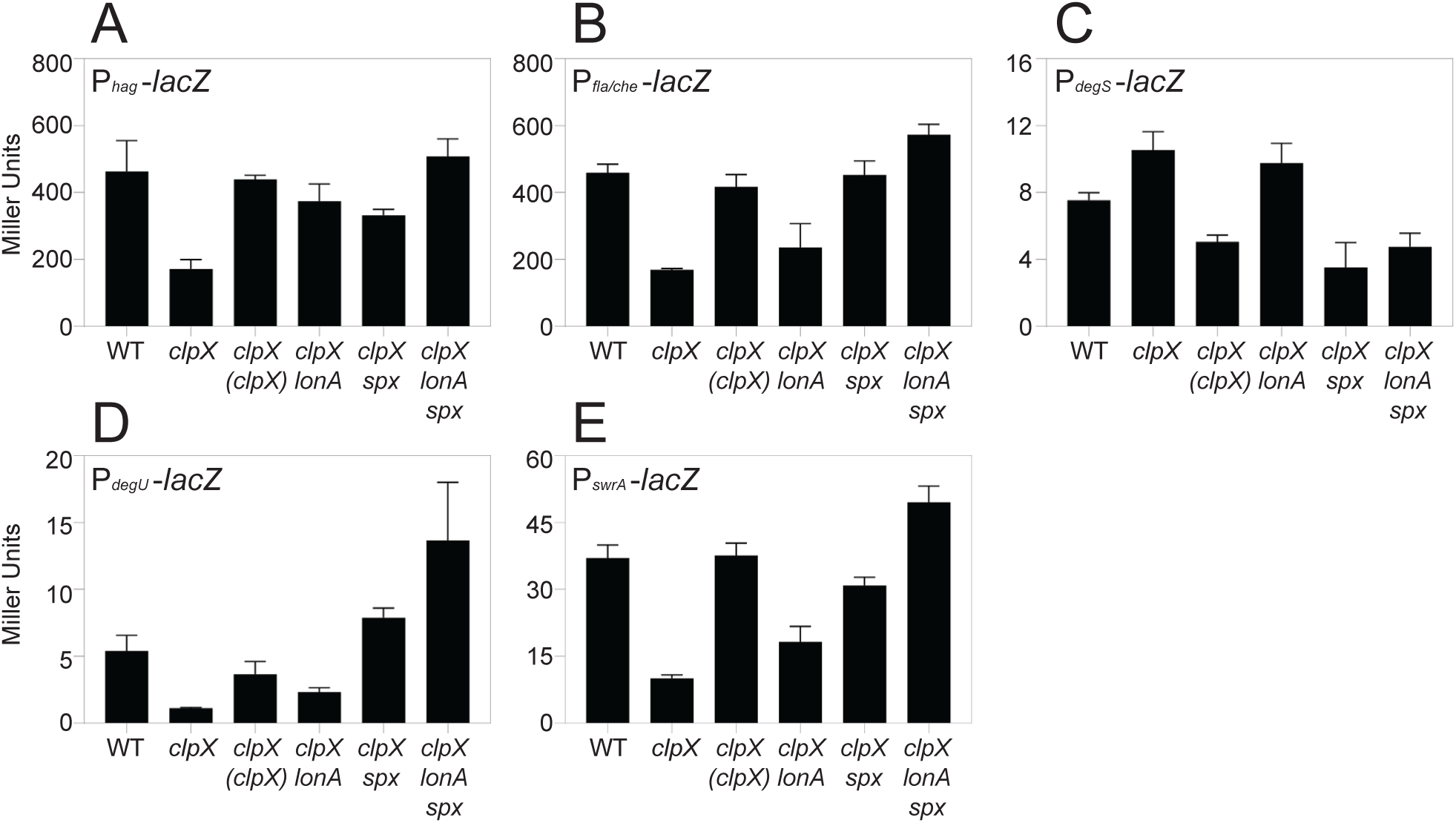
Mutation of ClpX reduces transcription from a variety of transcriptional reporters. β-galactosidase assays measuring transcriptional activity from key motility promoters in a variety of genetic backgrounds. Genes in italics are mutated, and genes in parentheses are complementation constructs expressed under the gene’s native promoter and inserted at an ectopic site in the chromosome. Each bar shows the mean Miller Units of three biological replicates with error bars indicating standard deviation. (A) Expression from the promoter of the *hag* gene (*P_hag_*-*lacZ*). The following strains were used to generate the data in this panel: WT (DK5457), *clpX* (DB20), *clpX* (*clpX)* (DB26), *clpX lonA* (DB28), *clpX spx* (DB24), *clpX lonA spx* (DB54). (B) Expression from the promoter of the *fla/che* operon (*P_fla/che_*-*lacZ*). The following strains were used to generate the data in this panel: WT (DK2399), *clpX* (DB67), *clpX* (*clpX)* (DB68), *clpX lonA* (DB70), *clpX spx* (DB69), *clpX lonA spx* (DB71). (C) Expression from the promoter of the *degSU* operon (*P_degS_*-*lacZ*). The following strains were used to generate the data in this panel: WT (DK9529), *clpX* (DB1527), *clpX* (*clpX)* (DB1529), *clpX lonA* (DB1530), *clpX spx* (DB1528), *clpX lonA spx* (DB1531). (D) Expression from the internal promoter directly upstream of the *degU* gene (*P_degU_*-*lacZ*). The following strains were used to generate the data in this panel: WT (DB559), *clpX* (DB807), *clpX* (*clpX)* (DB811), *clpX lonA* (DB813), *clpX spx* (DB809), *clpX lonA spx* (DB848). (E) Expression from the promoter of the *swrA* gene (*P_swrA_*-*lacZ*). The following strains were used to generate the data in this panel: WT (DK6613), *clpX* (DB806), *clpX* (*clpX)* (DB810), *clpX lonA* (DB812), *clpX spx* (DB808), *clpX lonA spx* (DB847).

The P*_hag_* promoter is expressed by RNA polymerase and the alternative sigma factor SigD (Mirel 1989), and thus a reduction in P*_hag_* expression could be due to a reduction in SigD activity (**Fig 1**). SigD activity is antagonized by direct binding with its cognate anti-sigma factor FlgM (Caramori 1996; Fredrick 1996; Bertero 1999). Reduced SigD activity alone however was insufficient to explain the swarming defect of the *clpX* mutant as simultaneous mutation of ClpX and FlgM did not restore swarming (**Fig S1A**). SigD activity is also controlled by the level of SigD protein, and not only were SigD levels slightly reduced in the *clpX* mutant, the levels increased to wild type when *clpX* was ectopically complemented (**Fig 5**). We conclude that although reduced SigD activity is not responsible for the lack of swarming in the *clpX* mutant, the reduction in SigD levels may indicate that the effect of ClpX is upstream of SigD expression.

One way in which ClpX could act upstream to modulate SigD levels is by increasing expression of the *sigD* gene. The gene encoding SigD, *sigD*, is the penultimate gene of the 27kb-long *fla/che* operon that encodes many of the proteins required for the assembly of the flagellar basal body (**Fig 1**) (Helmann 1988; Albertini 1991; Zuberi 1991; Marquez Magana 1994). To determine whether mutation of ClpX affects basal body formation, GFP was fused to the C-ring protein FliM that forms puncta indicative of intact basal bodies (Guttenplan 2013; Dunn 2025). Cells mutated for ClpX showed a reduction in the number of flagellar basal bodies relative to wild type, with a corresponding increase in the amount of cytoplasmic fluorescence presumably due to excess unassembled FliM (**Fig 4B**). Puncta number increased and cytoplasmic fluorescence decreased upon ectopic complementation of *clpX* (**Fig 4B**). We conclude that both SigD levels and the number flagellar basal bodies decreases in the absence of ClpX, both likely due to decreased expression of the *fla/che* operon.

Expression of the *fla/che* operon is controlled by the P*_fla/che_* promoter recognized by RNA polymerase and the vegetative sigma factor SigA (**Fig 1**) (Estacio 1998; West 2000). A reduction in transcription from P*_fla/che_* could explain both the reduction in basal body number and SigD levels. Expression from a transcriptional reporter in which P*_fla/che_* was fused to the *lacZ* gene was reduced two-fold from wild type levels in the absence of ClpX, a level of reduction previously shown to be sufficient to abolish swarming motility (Kearns 2005; Mukherjee 2015), (**Fig 6B**), and expression was restored by *clpX* ectopic complementation. To determine whether the inhibition of the *fla/che* operon was sufficient to abolish swarming motility, the native P*_fla/che_* promoter was replaced with an artificial IPTG-inducible promoter. In wild type cells motility was IPTG-dependent, and induction with IPTG resulted in rapid swarming without the typically observed lag period (**Fig 2B**). Induction of the *fla/che* operon in the absence of ClpX restored swarming, albeit with a lag period similar to that observed in wild type cells (**Fig 2B**). We conclude that overexpression of the *fla/che* operon is sufficient to restore swarming in the absence of ClpX. We infer that the inhibition of flagellar motility by ClpX occurs at or before the level of P*_fla/che_*, consistent with previous results and models (Moliere 2016).

### ClpX enhances swarming motility through the SwrA•DegU master activator

P*_fla/che_* is activated by a heteromeric transcription factor complex made of the response regulator DegU and the accessory protein SwrA (Tsukuhara 2008; Ogura 2012; Mordini 2013; Mishra 2024) (**Fig 1**), and Western blot analysis indicated that the levels of both DegU and SwrA were reduced in the absence of ClpX (**Fig 5**). The gene *degU*, encoding DegU, is expressed from its own promoter but is also part of a two-gene operon preceded by the gene *degS*, encoding DegS, the DegU-cognate histidine kinase (**Fig 1**) (Msadek 1990; Mukai 1990; Yasamura 2008; Shiwa 2015). Expression of a *lacZ* transcriptional reporter to P*_degS_* marginally increased compared to wild type in the absence of ClpX (**Fig 6C**), but expression decreased from reporters of *P_degU_* (**Fig 6D**) and *P_swrA_* (**Fig 6E**), following the trends previously observed with other motility promoters. Transcription from both P*_degU_* and P*_swrA_* could be restored by ectopic complementation of *clpX* (**Fig 6D**, **Fig 6E**). We conclude that the reduction of swarming motility in the absence of ClpX is correlated with a reduction in the transcription of genes encoding DegU and SwrA.

To determine if the repression of DegU and SwrA was sufficient to account for the swarming defect in the absence of ClpX, artificial overexpression constructs were introduced into wild type and ClpX mutant backgrounds. Overexpression of either *degU* or *swrA* abolished the swarming lag period in wild type cells and increased swarming motility in the absence of ClpX (**Fig 2C**, **Fig 2D**). Simultaneous overexpression of both SwrA and DegU also enhanced swarming of the *clpX* mutant and did so to a higher level than overexpression of either SwrA or DegU alone (**Fig 2E**). We conclude that the reduced expression of each component of the hybrid master regulator of flagellar biosynthesis contributes to the swarming defect of a *clpX* mutant.

### Mutations that enhance *fla/che* expression suppress the absence of ClpX

To determine how swarming is inhibited in the absence of ClpX, twenty-three spontaneous suppressors that restored swarming motility to a *clpX* mutant were clonally isolated and mapped by either candidate-gene or whole-genome sequencing. The suppressor mutations were organized into three genetic classes (**Table 1**). Class I suppressors were in or near the P*_fla/che_* promoter of the *fla/che* flagellar operon. One suppressor contained a mutation that changed the -10 box closer to consensus for SigA (Class Ia), and another suppressor deleted an intrinsic terminator of *codY*, the gene immediately upstream of the *fla/che* operon (Class Ib). Both mutations have been observed in other suppressor selection screens, and have been shown to increase *fla/che* operon expression (Amati 2004; Kearns 2005; Phillips 2015) (**Fig S2A, Fig S2B**). We conclude that both Class I mutations increase *fla/che* transcription, thereby bypassing the need for ClpX. We further conclude that the mutations support the inference that ClpX acts at or above the level of the P*_fla/che_* promoter.

Class II suppressors contained frameshift mutations in the gene *lonA*, encoding LonA, a protease responsible for the regulatory proteolysis of SwrA (**Fig S2C**). Mutation of *lonA* increased swimming motility in the absence of ClpX (**Fig 3**) and also partially restored the ability to swarm (**Fig 2F**). The improved motility of the *clpX lonA* double mutant was correlated both with an increase in the number of flagellar basal bodies (**Fig 4B**) and in the number of flagellar filaments on the cell surface (**Fig 4A**). Moreover, defects in Hag and SigD protein levels (**Fig 5**) as well as reductions in *hag* and *fla/che* reporter expression were increased in the double mutant (**Fig 6A, 6B**). The levels of *swrA* and *degU* expression by transcriptional reporter activity also increased in the *clpX lonA* double mutant (**Fig 6D**, **Fig 6E**), and while SwrA protein levels increased dramatically, DegU protein levels did not (**Fig 5**). We conclude that the absence of LonA improves each of the motility phenotypes tested in cells lacking ClpX by increasing SwrA protein levels, which in turn, increases *fla/che* operon transcription. Based on the Class I and II mutations, we conclude that one way to enhance swarming motility in the absence of ClpX is by increasing expression of the *fla/che* operon.

### Mutation of Spx suppresses the absence of ClpX by increasing SwrA and DegU levels

The final and largest class of suppressors was Class III that disrupted the gene *spx*, encoding the global transcriptional regulator Spx. Three of the Class III suppressor mutants contained large deletions in the *spx* open reading frame (Class IIIa) (**Fig S2D**), and another two suppressors contained even larger deletions of multiple genes that included *spx* (Class IIIb) (**Fig S2E**). Four suppressors were point mutations in the -10 box of the *spx* promoter (P*_spx_*) such that it was farther from consensus for SigA (Class IIIc) (**Fig S2F**), and one was a -10 box deletion (Class IIId). Finally, seven suppressors introduced missense mutations in the *spx* coding region (Class IIIe) (**Fig S2G-H**), while an eighth inserted one full codon into the reading frame (Class IIIf) (**Fig S2I**). Spx is degraded by ClpXP with the aid of the adaptor protein YjbH and is often isolated as a <u>s</u>uppressor of *clp<u>P</u>* and *clpX* mutant phenotypes (Nakano 2001; Larsson 2007; Garg 2009). In sum, we infer that Class III suppressor mutations likely decrease Spx levels and/or function.

To explore the mechanism of Spx-mediated repression of motility, we introduced a mutation in *spx* to our *clpX* mutant cells and assessed each of the phenotypes previously tested. Consistent with the spontaneous mutants, mutation of Spx increased swarming motility in the absence of ClpX (**Fig 2G**), and also enhanced swimming (**Fig 3**). Likewise, mutation in *spx* enhanced flagellin gene expression (**Fig 6A**), flagellin protein levels (**Fig 5**), basal body number (**Fig 4B**), and flagellar number per cell (**Fig 4A**) when ClpX was mutated. While Hag and SigD protein levels appeared to be increased to wild type levels in the *clpX spx* double mutant, the increase in the levels of both DegU and SwrA appeared to exceed that of the wild type (**Fig 5**). Consistent with elevated DegU and SwrA, we observed a restoration in expression from P*_fla/che_* in the *clpX spx* double mutant (**Fig 6B**). We conclude that Spx accumulation inhibits the accumulation of SwrA and DegU, and the reduction in the flagellar master regulator components restricts the cells’ ability to produce flagella sufficient for swarming.

To determine whether Spx inhibited SwrA and DegU expression at the transcriptional level, reporter gene expression assays were conducted. Expression from the P*_degU_* and P*_swrA_* reporters was increased in the *clpX* mutant background when *spx* was also mutated (**Fig 6D-E**), but expression from the P*_degS_* promoter was not (**Fig 6C**). We conclude that in cells mutated for ClpX, Spx accumulates and most likely represses motility by inhibiting transcription from P*_degU_* and P*_swrA_* as artificial induction of both *swrA* and *degU* restored swarming to our *clpX* mutant close to wild-type levels (**Fig 2E**). Thus, Spx inhibits motility at the highest level by restricting expression of the master activator components.

Our genetic analysis suggests that swarming motility can be restored to a *clpX* mutant by increasing expression of the *fla/che* operon, either directly by improving the promoter, or by increasing accumulation of SwrA and DegU. Mutation of LonA restores swarming by relieving proteolytic restriction of SwrA levels, and mutation of Spx restores swarming by relieving repression on both *degU* and *swrA* gene expression. Thus, each suppressor seems to restores components of the hybrid master regulator in different ways. Consistent with parallel mechanisms, cells mutated for ClpX, LonA, and Spx simultaneously increased swarming (**Fig 2H**), P*_fla/che_*, P*_degU_* and P*_swrA_* reporter gene expression (**Fig 6B, 6D, 6E**), and SwrA protein (**Fig 5**) to a levels greater than either *clpX* containing double mutant alone. We conclude that Spx and LonA act in parallel and by separate mechanisms to restrict flagellar biosynthesis at level of the master activator of flagellar biosynthesis.

## DISCUSSION

Flagellar gene expression is hierarchical and is organized in at least three regulatory tiers (Dasgupta 2003; Chevance 2008; Smith 2009; Ardissone 2015;). In *Bacillus subtilis*, the first tier is the hybrid master activator SwrA•DegU, which in turn activates the second tier *fla/che* operon encoding the basal body, rod, and hook structural components (Mukherjee 2014). The third tier constitutes a regulon of genes under control of the alternative sigma factor SigD (Serizawa 2004; Kearns 2005), which becomes activated after basal body-rod-hook assembly is complete (Barilla 1994; Calvo 2015), and contains genes required for the subsequent assembly of the flagellar filament (Mirel 1989; Chen 1994). Previous work indicated that when ClpX is mutated, Spx accumulates and inhibits swimming motility at the level of *fla/che* operon expression (tier II) (Moliere 2016; Schafer 2019). Here, we show that swarming motility is also inhibited in the absence of ClpX, and genetic suppressor analysis indicated that Spx inhibits the master activator (tier I). Thus, our work reinforces previous observations that Spx inhibits expression of the *fla/che* operon, but does so at one level higher than previously recognized by restricting production of both SwrA and DegU.

Spx is an unusual transcription factor in that it acts as a conditional subunit of RNA polymerase, binding between one of the alpha subunits and the vegetative sigma factor SigA (Birch 2017; Shi 2021). Spx is thought to activate the expression of specific genes by stabilizing RNA polymerase interaction at promoters that contain a particular -44 element (Reyes 2008; Nakano MM 2010 Rochat 2012). Spx is thought to repress gene expression by a similar mechanism, whereby its presence in RNA polymerase interferes with transcriptional activators that require contact with the alpha subunit’s C-terminal domain (Nakano S 2003; Newberry 2005; Zhang 2006). Here we show that high levels of Spx inhibit transcription from the P*_swrA_*, P*_degU_*, and P*_fla/che_* promoters, and the effect can be bypassed either by artificial expression of the *fla/che* operon or both SwrA and DegU together. Consistent with sequence-independent inhibition, a previously published ChIP seq analysis of Spx binding sites, indicates that Spx does not bind to any of these promoters (Rochat 2012). Thus, our results seem consistent with a model whereby Spx inhibits by a DNA-independent anti-alpha mechanism, and it may antagonize the expression of all three motility promoters in parallel, as they are all activated by the transcription factor DegU (Cairns 2013; Ogura 2010; Tsukahara 2008; Mishra 2024).

DegU is a highly-pleiotropic DNA-binding response regulator, but the sequence to which it binds and the mechanism of transcriptional activation are poorly understood. Recent work indicated that DegU binds upstream of the P*_fla/che_* promoter but does not activate it until SwrA interaction causes DegU to oligomerize, perhaps facilitating interaction with the alpha C-terminal domain of RNA polymerase (Tsukahara 2008; Ogura 2012; Mordini 2013; Mishra 2024). ChIP-seq analysis also indicated that DegU bound near the P*_swrA_* and P*_degU_* promoters *in vivo*, and DegU has been shown to be required for the activation of each (Calvio 2008; Ogura 2010; Cairns 2013; Mishra 2024). As DegU activates all of the promoters that appear to be antagonized by Spx, we infer that at high levels Spx inhibits swarming motility by blocking SwrA•DegU autoactivation. Consistent with Spx inhibiting both components of the master activator, mutation of Spx bypasses the need for ClpX and restores the levels of both proteins and robust swarming (**Fig 2G**), whereas mutation of LonA partially compensates by increasing SwrA alone, and swarming restoration is weak (**Fig 2F**). We infer that the primary role of Spx is to inhibit DegU-dependent activation, as high Spx levels inhibit swimming in laboratory strains that naturally lack SwrA (Moliere 2016; Kearns 2004; Calvio 2005; Gallegos-Monterrossa 2016), and DegU activates other promoters related to motility (Hsueh 2011).

Spx is normally kept at a low level in the cell by ClpXP and the proteolytic adaptor YjbH (Larsson 2007; Garg 2009), and proteolysis is thought to be relieved under stress conditions (Petersohn 2001; Nakano S 2003; Nakano S 2005). Why flagellar motility would be inhibited under stress is unclear, and it is also unclear which stress conditions would lead to Spx-mediated inhibition. One possible stress could be flagellar assembly itself; when cells are placed on a swarming-permissive surface, the density of flagella on the cell surface doubles and reaches nearly 30 flagella per cell (Mukherjee 2015). How flagella are assembled through the *B. subtilis* cell wall is unknown but could require peptidoglycan remodeling and/or insertion during peptidoglycan synthesis (Mukherjee 2014; Sanchez 2021; Sanchez 2022; Dunn 2025). In any case, insertion of a large number of transenvelope machines may trigger a peptidoglycan stress response, and recent work has indicated that cell wall stress causes Spx levels to rise (Rojas-Tapias 2018). Our model suggests that Spx may dampen a DegU-mediated positive feedback loop, and Spx could be a means to restrict cell wall damage from runaway flagellar biosynthesis.

While our data largely supports a mechanism in which Spx antagonizes DegU-dependent transcriptional activation, we note that two observations seem inconsistent with the generalized anti-alpha model of Spx repression. First, overexpression of both SwrA and DegU was sufficient to override high levels of Spx (**Fig 2E**). If Spx indeed acts to make RNA polymerase unreceptive to the presence of transcriptional activators, it seems inconsistent that high levels of an activator would simply override its presence. Second, the model that Spx acts like an anti-alpha factor is partly predicated on its inhibition of the ComA-activated P*_srfAA_* promoter in domesticated laboratory strains that do not produce surfactin (Nakano MM 1992; Nakano S 2003; Lin 2013). However, we find that the inhibition of swarming motility by high levels of Spx in the ancestral strain is not due to a lack of surfactin production. Indeed, the *clpX* mutant produced levels of surfactin that were not only detectable to the naked eye but also sufficient to select for suppressors that restore swarming. Why Spx fails to restrict surfactin production under swarming conditions is unclear, but perhaps the inhibitory activity of Spx is more specific than previously recognized. Precisely how Spx antagonizes transcription remains unknown.

## ACKNOWLEDGEMENTS

We thank the Indiana University Center for Genomics and Bioinformatics for whole genome sequencing and subsequent data analysis. This work was funded by NIH grant R35 GM131783 to DBK.

---

**Figure S1. Mutation of the anti-sigma factor FlgM does not restore swarming in cells mutated for ClpX.**

Quantitative swarm expansion assays for the strains indicated in each panel. Genes in italics are mutated, genes in parenthesis are complementation constructs expressed under the gene’s native promoter and inserted at an ectopic site in the chromosome, and a “+” indicates that the gene was induced with 1 mM IPTG. Each data point is the average of three technical replicates. (A) The *clpX* mutant phenotype was not rescued when the gene for the anti-sigma factor, *flgM*, was also mutated. The following strains were used to generate the data in this panel: WT (DK1042), *clpX* (DK6563), *clpX flgM* (DB3069). **(B)** The partially rescued *clpX degU*^+^ phenotype (see Figure 2C) was not improved when the gene for the anti-sigma factor, *flgM*, was also mutated. The following strains were used to generate the data in this panel: WT (DK1042), *clpX degU*^+^ *flgM* (DB3070, 1 mM IPTG added), *clpX* (DK6563).

**Figure S2. Spontaneous suppressor mutations restore swarming in cells mutated for ClpX.**

Quantitative swarm expansion assays for the strains indicated in each panel. “*sox*” or “*axs*” indicate suppressor of *clpX* isolated from separate experiments. Genes in italics are mutated. Each data point is the average of three technical replicates. (A) The *clpX* mutant phenotype was rescued by a single nucleotide polymorphism in P*_fla/che_* (*sox16*). The following strains were used to generate the data in this panel: WT (DK1042), *sox16* (DK9447), *clpX* (DK6563). (B) The *clpX* mutant phenotype was rescued by a 25-nucleotide deletion including the terminator for *codY* (*sox15*). The following strains were used to generate the data in this panel: *sox15* (DK9750), WT (DK1042), *clpX* (DK6563). (C) The *clpX* mutant phenotype was rescued by mutations in *lonA*: *sox9* and *sox12* were slipped-strand mispairing mutations and *sox17* was an insertion of 23 nucleotides. The following strains were used to generate the data in this panel: WT (DK1042), *sox9* (DK9748), *sox17* (DK9751), *sox12* (DK9749), *clpX* (DK6563). (D) The *clpX* mutant phenotype was rescued by deletions in the *spx* coding region: *sox2* was a deletion of coding nucleotides, and *sox18* and *sox19* were deletions including the promoter and coding nucleotides. The following strains were used to generate the data in this panel: WT (DK1042), *sox2* (DK9386), *sox19* (DK9391), *sox18* (DK9390), *clpX* (DK6563). (E) The *clpX* mutant phenotype was rescued by different large deletions of multiple genes that included *spx* (*sox1* and *sox6*). The following strains were used to generate the data in this panel: WT (DK1042), *sox6* (DK9746), *sox1* (DK9744), *clpX* (DK6563). (F) The *clpX* mutant phenotype was rescued by a loss-of-function SNP in P*_spx_* (*sox5*, *sox10*, *sox11*, and *sox13*). Only *sox5* is shown. The following strains were used to generate the data in this panel: WT (DK1042), *sox5* (DK9393), *clpX* (DK6563). (G) The *clpX* mutant phenotype was rescued by missense mutations in the *spx* coding region: *sox14* = *spx*^R119H^, *sox3* = *spx*^F113C^, *sox20* = *spx*^V2F^, *sox7* = *spx*^Y5C^. The following strains were used to generate the data in these panels: WT (DK1042), *sox14* (DK9389), *sox3* (DK9387), *sox20* (DK9392), *sox7* (DK9388), *clpX* (DK6563). (H) The *clpX* mutant phenotype was rescued by more missense mutations in the *spx* coding region: *asx27* = *spx*^T49I^, *asx17* = *spx*^P93L^, *asx6* = *spx*^Y80H^. Suppressors are shown in two separate panels for simplicity. The following strains were used to generate the data in these panels: WT (DK1042), *asx27* (DK9865), *asx17* (DK9843), *asx6* (DK9753), *clpX* (DK6563). (I) The *clpX* mutant phenotype was rescued by an inserted glycine at amino acid position 53 in *spx* (*asx4*). The following strains were used to generate the data in this panel: WT (DK1042), *asx4* (DK9842), *clpX* (DK6563). See Table 1 for more detailed information on suppressor mutations.

